# Impaired proteasome induces mitochondrial DNA release to activate the cGAS-STING signaling pathway and cause necroptosis in mouse brain

**DOI:** 10.1101/2024.11.13.623442

**Authors:** Abena Dwamena, Yasin Asadi, Erin Gilstrap, Hongmin Wang

## Abstract

Impaired proteasome function is consistently associated with many neurodegenerative disorders, including Alzheimer’s disease (AD), showing neuroinflammation and neurodegeneration; however, how impaired proteasome causes neuroinflammation and neuronal death remains less understood. Here, we studied the effect of impaired proteasome on neuroinflammation and neuronal death in a knockout (KO) mouse model with reduced proteasome activity in the brain. We discovered that impaired proteasome led to the release of mitochondrial dsDNA into the cytosol, activating the cGAS-STING signaling pathway, and upregulating pro-inflammatory cytokines in the KO mouse brain relative to the control brain. Importantly, we also observed reduced brain weight, elevation of the mixed lineage kinase domain-like (MLKL) protein, phosphorylated MLKL, and receptor-interactive protein kinases (RIPK) 1 and 3 in the KO mouse brain compared to the control brain, suggesting activation of necroptosis in the KO brains. These data indicate that impaired proteasome activates the cGAS-STING pathway to induce neuroinflammation and neurodegeneration via a necroptotic manner. Our results suggest that neuroinflammation and necroptosis may be generalized factors caused by reduced proteasome activity observed in diverse neurodegenerative disorders.

## Introduction

As a major intracellular protein degradation system, the proteasome is responsible for the turnover of at least 80% of total proteins in many types of cells (Collins & Goldberg, 2017). However, proteasome function is impaired in the condition of aging and many neurodegenerative disorders, including Alzheimer’s disease (AD) (Wang & Wang, 2020). Co-occurrence with impeded proteasome is mitochondrial fragmentation and neuroinflammation in the brain affected by numerous neurodegenerative diseases (Guzman-Martinez *et al*, 2019; Huber *et al*, 2024; Ugun-Klusek *et al*, 2017). How impaired proteasome is linked to neuroinflammation and neuronal death in the brain remains unknown.

The Cyclic GMP-AMP synthase (cGAS) is a major cytoplasmic DNA receptor recognizing both endogenous and exogenous double-stranded DNA (dsDNA). Upon binding to dsDNA, cGAS catalyzes a biochemical reaction using both guanosine triphosphate (GTP) and adenosine triphosphate (ATP) to generate a secondary messenger, known as the cyclic GMP-AMP (2′3′-cGAMP) that functions as an endogenous ligand to bind to and activate the key adaptor protein, stimulator of interferon gene (STING) (Gulen *et al*, 2023; Yu & Liu, 2021). Activation of the cGAS-STING pathway in various pathological conditions, including tauopathy/AD, induces neuroinflammation by upregulating the downstream target genes such as NF-κB, TNF-α, IL1b, and IL-6 (Udeochu *et al*, 2023; Zhang & Zhong, 2022).

Necroptosis, a programmed form of necrosis, has been implicated in aging and many neurological diseases. Necroptosis is primarily triggered by the activation of the Mixed Lineage Kinase Domain-Like (MLKL) and Receptor-Interacting Protein Kinases (RIPK) 1 and 3. Essentially, when RIPK1 and RIPK3 are activated, they phosphorylate MLKL (pMLKL), resulting in its *oligomerization* and eventually disrupting the cell membrane and causing necroptotic cell death (Galluzzi *et al*, 2017).

Because AD and many other aging-related neurodegenerative disorders are characterized by impaired proteasome, neuroinflammation, and loss of neurons, we here to test whether 26S proteasome deficiency causes neuroinflammation and cell death via release of mitochondrial (mt) DNA and activation of cGAS-STING pathway in the brain. We took advantage of a conditional knockout mouse model to genetically ablate *Psmc1*, one 19S proteasomal subunit of the 26S proteasome, in neurons of the forebrain region (Bedford *et al*, 2008; Huber *et al*., 2024; Ugun-Klusek *et al*., 2017). With this model, we found that impaired 26S proteasome resulted in increased cytosolic deposit of mitochondrial dsDNA, activation of the cGAS-STING pathway, neuroinflammation, and necroptosis.

## Results

### Impaired proteasome induces cytosolic dsDNA deposits of mitochondrial origin in the brain cortex

We recently confirmed proteasome dysfunction in the brain of a mouse model deficient in the 26S proteasome, where one 19S proteasomal subunit, *Psmc1*, is genetically ablated selectively in the forebrain region (Huber *et al*., 2024). Because a deficiency of Psmc1 (KO) was previously shown to cause marked mitochondrial fragmentation in the brain (Ugun-Klusek *et al*., 2017), we, therefore, determined whether the cytosolic release of the double-strand (ds) DNA could be detected in the brain by an immunohistochemical approach using an antibody specifically recognizing *cytosolic* dsDNA but not the highly packed nuclear DNA (Ding *et al*, 2018). As demonstrated in **Fig. 1A** and **1B**, KO of *Psmc1* at three months of age increased cytosolic dsDNA staining in the brain cortex compared to the control animal brains. To further determine the origin of dsDNA, we performed subcellular fractionation to isolate the cytosol after removing the mitochondrial and nuclear fractions of the brain tissue cells. PCR amplification of the cytoplasmic mitochondrial DNA (mtDNA), using the primers selectively amplifying the cytochrome c oxidase 1 (COX1) gene, showed a significantly increased level of mtDNA in the KO mice compared to the control (**Fig. 1C**). These data suggest that when proteasome function is disrupted, it leads to the deposition of mitochondrial DNA in the cytosol.

**Fig. 1.**
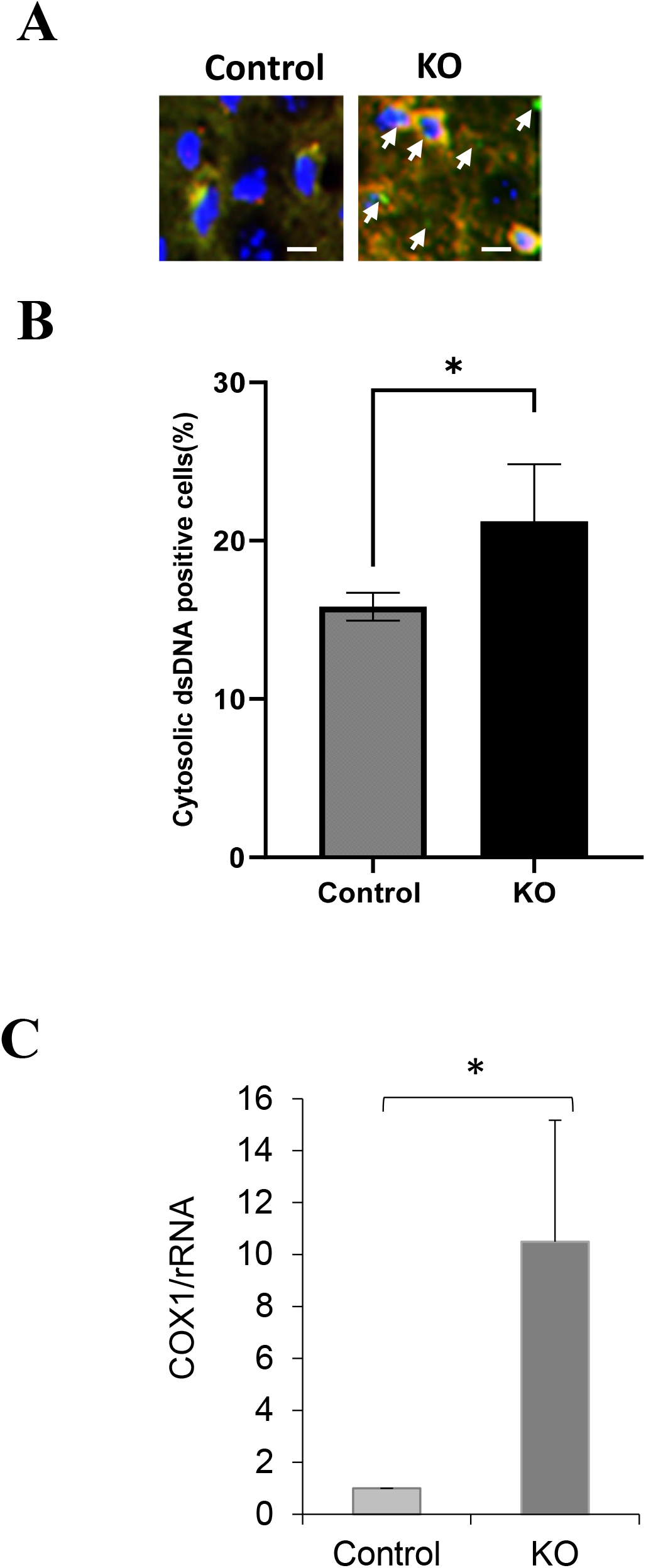
Impaired proteasome induces cytosolic dsDNA deposits of mitochondrial origin in the brain cortex. **A**. Immunofluorescent staining of cytosolic dsDNA in the brain cortex at 3 months of age. Red, mitochondria; Green, dsDNA. Scale bar, 10 μm. **B**. Quantitation of cytosolic dsDNA staining in the brain cortex. **C**. qPCR analysis of cytosolic mtDNA. Data are presented as mean ± SD; n = 6, * p < 0.05.

### Impaired proteasome activates the cGAS-STING signaling pathway

As we observed the presence of cytosolic mtDNA, we went on to test whether this activates a cellular DNA-sensing pathway, the cGAS-STING signaling. The cGAS-STING signaling pathway is a part of the innate immune system that detects the presence of DNA in the cytosol to activate the pro-inflammatory genes, such as p-IRF3, NF-κB, and their downstream targeting genes (**Fig. 2A**). We therefore examined the proteins involved in this pathway at three months with Western blot analysis, including cGAS, STING, p-TBK, TBK, IRF3, and p-IRF3. As shown in **Fig. 2B and 2C**, all the examined proteins showed marked increases in the cortex of the KO mouse brains when compared to the control mouse brains, depicting the activation of the cGAS-STING signaling pathway in the presence of mtDNA in the cytosol.

**Fig. 2.**
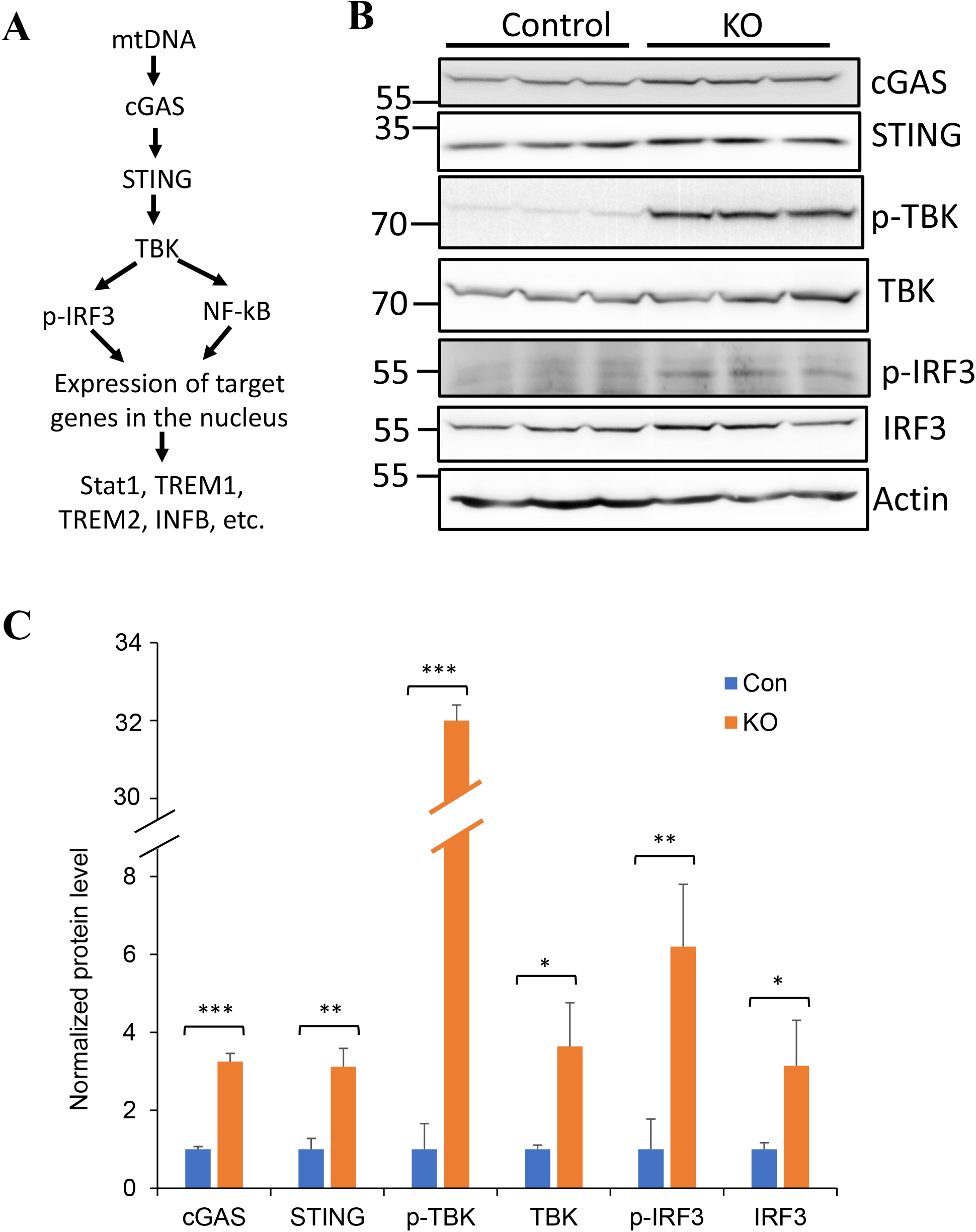
Impaired proteasome activates cGAS-STING signaling pathway. **A**. Scheme of cGAS-STING pathway activation upon mtDNA sensing in the cytosol. **B**. Western blot analysis of the indicated proteins involved in cGAS-STING pathway in the brain cortex at 3 months of age. **C**. Quantitation of cGAS-STING pathway protein levels. Data are presented as mean ± SD; n = 6, * p < 0.05, ** p < 0.01, *** p < 0.001.

### Impaired proteasome induces neuroinflammation

Since activation of the cGAS-STING pathway upregulates several key neuroinflammation modulators, such as NF-κB, Stat1, and TREM2 (**Fig. 2A**), we next assessed the level of these proteins at three months of age. Western blot analysis showed a significant elevation of NF-κB proteins in the KO mouse brain cortex compared to the control brain cortex (**Fig. 3A, 3B**). In addition to Stat1 and TREM2 proteins, our data also showed that some innate inflammation cytokines, such as IL1b, IL6, and TNFα, also showed marked increases in the KO brains compared to the control (**Fig. 3C, 3D**), suggesting activation of neuroinflammation in the KO brain.

**Fig. 3.**
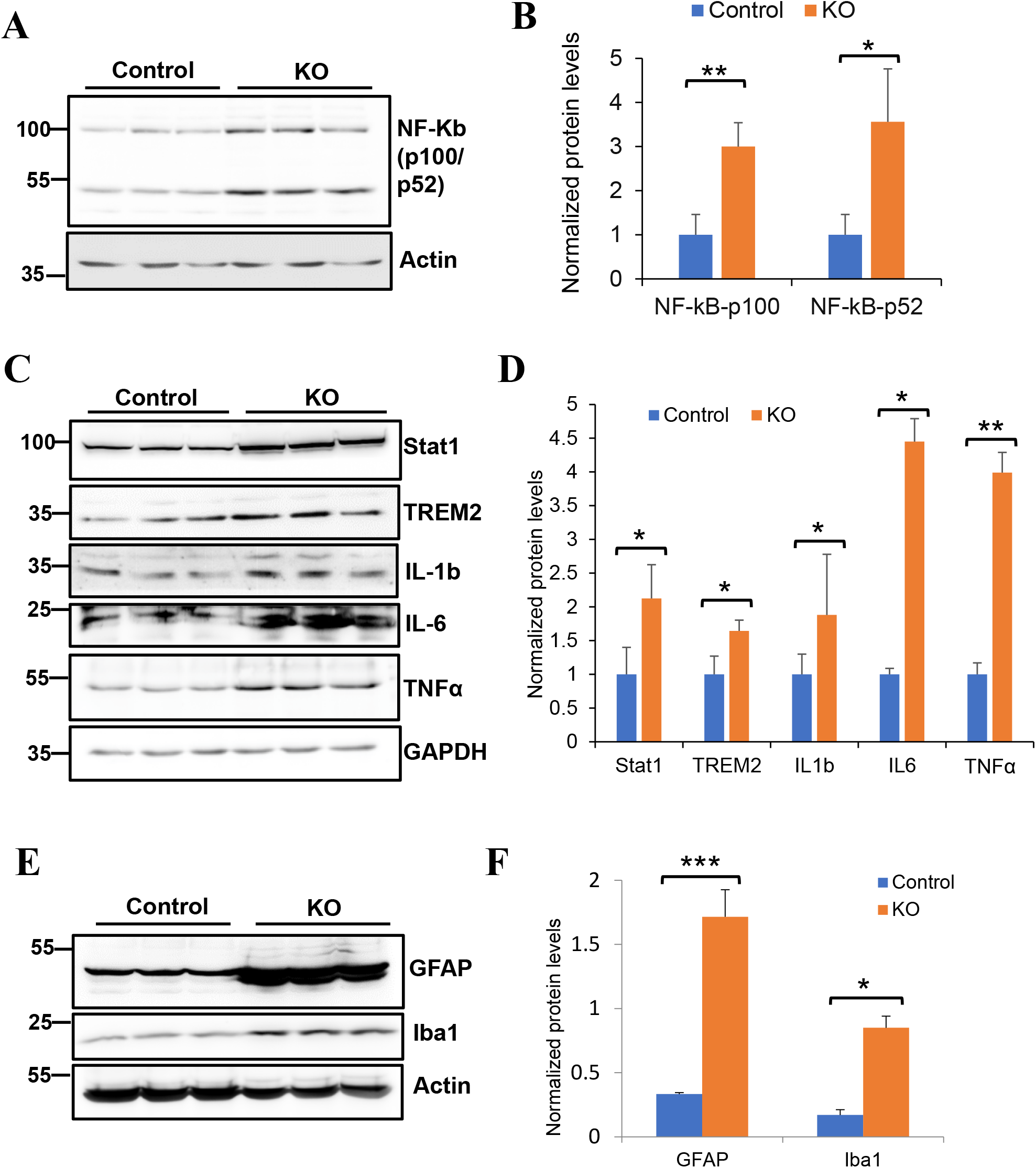
Impaired proteasome activates neuroinflammation and gliosis. **A**. Western blot analysis of NF-κB. It is noted that the two NF-κB members, p100 and p52, show marked upregulation in the KO brains compared to the control brains at 3 months of age. **B**. Quantitation of NF-κB protein. **C**. Western blot analysis of the indicated pro-neuroinflammatory cytokines. **D**. Quantitation of pro-neuroinflammatory cytokines. **E**. Western blot analysis of GFAP and Iba1 proteins. **F**. Quantitation of GFAP and Iba1 proteins. Data are presented as mean ± SD; n = 6. * p < 0.05; ** p < 0.01, *** p < 0.001.

Activation of astrocytes and microglia is a common feature of neuroinflammation and has been seen in many neurodegenerative diseases (Kwon & Koh, 2020). To assess whether the two types of glia were activated in the KO mouse brain at this age, we examined the level of the glial fibrillary acidic protein (GFAP), a protein indicative of astrocytes that represents a biomarker of neuroinflammation, and the ionized calcium-binding adaptor molecule 1 (Iba1), a marker for microglia, the resident macrophage cell in the brain (Guan *et al*, 2022), because upregulation of these proteins reflects an increased reactive phenotype (Wilhelmsson *et al*, 2006). Western blot analysis of GFAP and Iba1 indicated a significant increase of these proteins in the KO brain samples compared to controls (**Fig. 3E, 3F**). Taken together, these data strongly support that reduced proteasome activity in the KO mouse brain causes neuroinflammation and activation of astrocytes and microglia.

### Impaired proteasome reduces brain size

We isolated the brains and measured their weights to determine whether impaired proteasome activity causes neuronal loss and alters brain volume. As shown in **Fig. 4A**, the KO mouse brain appeared smaller than the control mouse brains at three months of age. The brain weight of the KO mice was lighter than that of the control mice (**Fig. 4B**), suggesting possible neurodegeneration in the KO mouse brains.

**Fig. 4.**
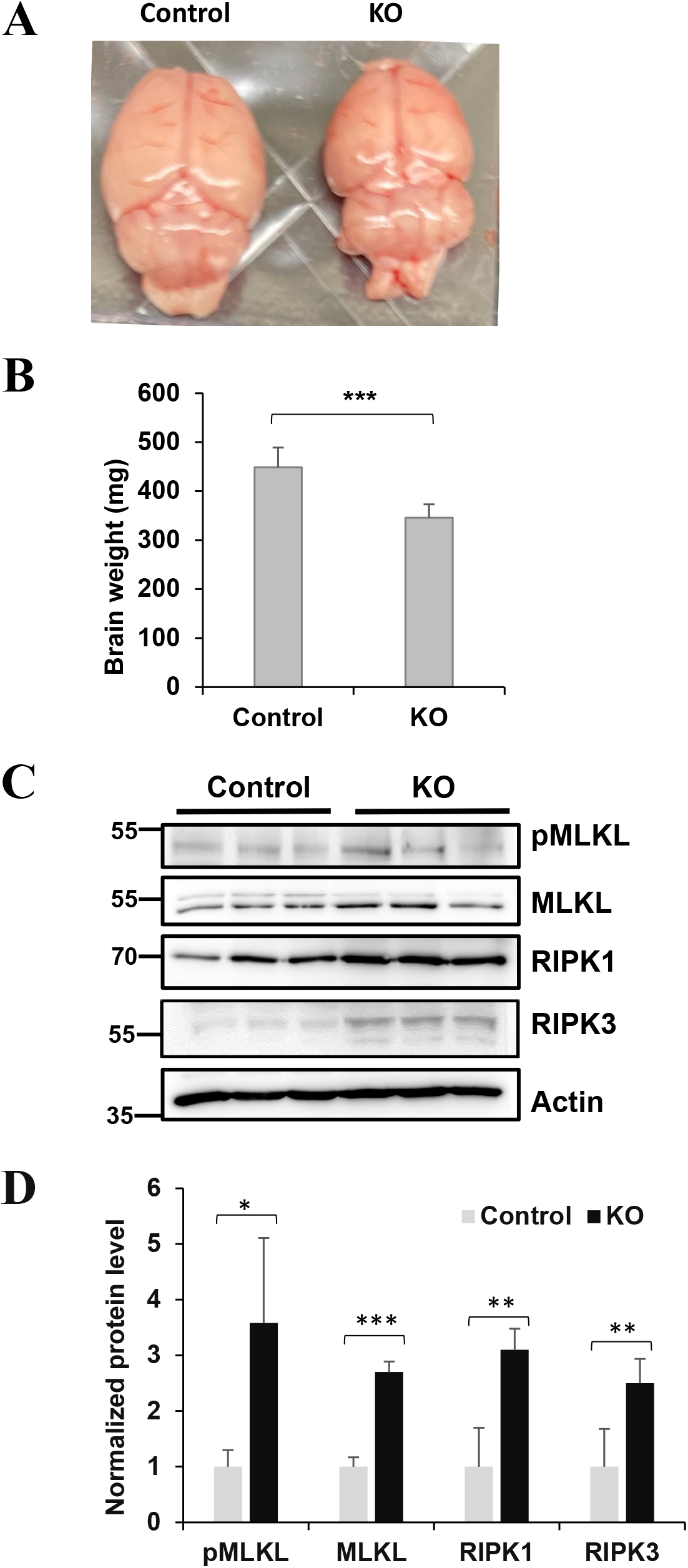
Impaired proteasome reduces brain volume and weight and activates necroptosis in *Psmc1* KO mouse brain. **A**. A KO brain appears smaller than a control brain. **B**. Reduce brain weight of KO brain compared to the control brain. **C**. Western blot analysis of the necroptotic proteins. **D**. Quantitation of necroptotic proteins. Data are presented as mean ± SD; n = 6. * p < 0.05; ** p < 0.01, *** p < 0.001.

### Impaired proteasome activates necroptosis

Apoptosis is a mechanism by which brain cells die in a programmed and controlled manner (Fricker *et al*, 2018). To determine whether the reduced brain volume and weight in the KO mice are due to increased apoptosis, we examined the level of cleaved caspase 3 at three months of age, a protein that is activated during apoptotic cell death and is commonly used as a reliable marker for cells that are dying or have died via apoptosis (Crowley & Waterhouse,2016). However, our Western blot results showed an undetectable cleaved caspase 3 from the KO mouse brains, similar to the control brains (data not shown), indicating the absence of overt apoptosis in the brain at this age.

Because necroptosis, a programmed form of necrosis, is closely associated with neuroinflammation (Caccamo *et al*, 2017), we next determined whether necroptosis was activated in the KO mouse brains where proteasome activity is reduced and neuroinflammation occurs (Huber *et al*., 2024). We performed Western blot analysis of the levels of the proteins executing necroptosis, RIPK1, RIPK3, MLKL, and pMLKL, in the brain cortex using buffers of increasing stringency (Caccamo *et al*., 2017). These proteins did not show a statistically significant difference between Control and KO brains in the Tris-buffered saline (TBS) fraction, or TBS containing 1% Triton X-100 (data not shown). In contrast, p-MLKL, MLKL, RIPK1, and RIPK3 protein levels in urea fractions were significantly higher in the KO brains than in the control brains (**Fig. 4C, 4D**). These results are consistent with previous observations that necroptotic markers, once activated, form *insoluble* amyloid-like structures (Caccamo *et al*., 2017).

## Discussion

Impaired proteasome functionality is implicated in many neurodegenerative disorders and is associated with neuroinflammation and neuronal death; however, how these occur remains evasive. In the mouse model with reduced proteasome activity, we not only confirmed earlier observations of neuroinflammation (Huber *et al*., 2024), but also expanded upon the findings by elucidating the involvement of the cGAS-STING pathway and necroptosis in the KO mouse brain.

Previous studies with the same mouse model indicated mitochondrial fragmentation and altered redox homeostasis in the mouse brain (Ugun-Klusek *et al*., 2017). Here, we demonstrated a significant increase in cytosolic release of mtDNA, which is supported by the immunofluorescence staining and high-sensitive qPCR. Unlike the hypermethylated nuclear DNA, mtDNA is hypomethylated, similar to bacterial DNA, which can potently activate innate immune signaling via the cGAS-STING pathway (Riley & Tait, 2020). We observed increased levels of cGAS, STING, and the downstream mediators such as p-TBK and p-IRF3, indicating robust activation of this pathway in the KO brain, which is further confirmed by the upregulation of the pro-inflammatory cytokines downstream of this pathway, such as Stat 1, IL1b, IL6, and TNFα, as well as by the increased GFAP and Iba1 levels.

In addition to apoptosis, brain neurons can also die through necroptosis, a manner of regulated necrosis cell death in various neurodegenerative diseases, including AD (Shao *et al*, 2022). Prior studies have indicated that neuronal necroptosis can occur in an apoptotic deficient condition (Naito *et al*, 2020). In the KO brain, we did not observe overt activation of caspase 3, suggesting the absence of apoptosis. Necroptosis and neuroinflammation appear to enhance each other. On one hand, necroptosis increases with aging in the brain and contributes to age-related neuroinflammation (Thadathil *et al*, 2021). On the other hand, necroptosis was originally identified as an inflammation condition and has been recognized as a cause of age-dependent neuroinflammatory diseases, including Alzheimer’s disease and Parkinson’s disease (Smith, 2024). Necroptosis is mediated by the RIPK1, RIPK3, and MLKL (Shan *et al*, 2018).

Importantly, both RIPK1 and RIPK3, as well as the p-MLKL (the active form of MLKL that causes the permeabilization of the plasma membrane, a crucial step in necroptosis) all showed elevated levels in the KO mouse brain, indicative of necroptosis.

In conclusion, this study reveals that impaired 26S proteasome causes the cytosolic release of mtDNA to activate the cGAS-STING signaling pathway, neuroinflammation, and necroptotic cell death in the brain.

## Materials and Methods

### Animals

All animal-related experiments and procedures were approved by the Institutional Animal Care and Use Committee at the Texas Tech University Health Science Center and compliant with the National Institute of Health Guide for the Care and Use of Laboratory Animals. Floxed *Psmc1* animals were described previously (Bedford *et al*., 2008; Huber *et al*., 2024; Ugun-Klusek *et al*., 2017). CamKIIα-Cre (T29-Cre) transgenic animals were acquired from the Jackson Laboratory (# 005359). The breeding strategy to produce the conditional *Psmc1* KO (fl/fl)/T29-Cre animals in the forebrain was based on the recommendation by the Jackson Laboratory.

### Detection of cytosolic dsDNA in brain sections with immunofluorescence

The immunofluorescence assay was based on a previous report (Liu *et al*, 2021). Briefly, after brain sections were permeabilized, they were incubated with a dsDNA primary antibody (1:50 dilution, Santa Cruz Biotechnology, #sc-58749) specifically recognizing mitochondrial (mt) DNA but not the highly packed nuclear DNA (Ding *et al*., 2018), and an anti-Cox-IV antibody (1:200 dilution, Cell Signaling Technology, #4850) for overnight at 4°C. Secondary antibody incubation, nuclear staining, and mounting slides were previously described (Liu *et al*, 2017). Stained brain sections were imaged and quantified with the ImageJ software as reported (Liu *et al*., 2021).

### Detection of cytosolic mtDNA with PCR

The cytosolic mtDNA was detected with the quantitative PCR (qPCR) as previously described (Guo *et al*, 2020). The primers for mt-Cox1 were 5′-GCCCCCGATATGGCGTTT-3′ (forward) and 5′-GTTCAACCTGTTCCTGCTCC-3′ (reverse). The primers for 18S rDNA were 5′-AGAGGGACAAGTGGCGTTC-3′ (forward) and 5′-CGCTGAGCCAGTCAGTGT-3′(reverse).

### Protein extraction

We followed the reported approach to extract the necroptosis-related proteins, pMLKL, MLKL, RIPK1, RIPK2 (Caccamo *et al*., 2017) to isolate the Tris-buffered saline (TBS), detergent-soluble (TBS containing 1% Triton X-100), and detergent-insoluble fractions. The detergent-insoluble fractions were re-suspended in 8 M urea containing 5% SDS and stored as urea fractions.

### Western blot analysis

Western blot analysis of brain proteins was based on previously described methods (Liu *et al*, 2020; Liu *et al*., 2021). Except anti-Actin (1:1000, Developmental Studies Hybridoma Bank), all other primary antibodies (1:1000) used here were from the Cell Signaling Technology. Protein band intensities were quantified using UN-Scan-IT Digitizer software (Silk Scientific).

### Statistical analysis

All numerical data are presented as mean ± SD. Student’s t-test was utilized to compare two groups. Statistical significance was accepted when p < 0.05.

## Acknowledgments

We would like to thank Drs. R. John Mayer and Zubair Karim for providing us with the floxed *Psmc1* mouse model. This work was supported in part by the NIH/NIA RF1 AG072510. Any opinions, findings, conclusions, or recommendations expressed in this material are those of the authors and do not necessarily reflect the views of the NIH/NIA.

## Author contributions

HW conceived the presented idea. AD, YA, EG, and HW designed and performed experiments. AD, YA, and HW analyzed the results. AD and HW interpreted the data and wrote the manuscript.

## Data availability

All data supporting the findings of this manuscript are available from the corresponding author upon reasonable request.

